# Fast analyzing, exploring and sharing quantitative omics data using omicsViewer

**DOI:** 10.1101/2022.03.10.483845

**Authors:** Chen Meng

## Abstract

**Summary:** To facilitate the biological interpretation of omics data, we developed a general omics data visualization platform termed omicsViewer, enabling interactive visualization of statistical results and performing downstream analyses directly in response to features and samples selected. Multiple hypotheses and parameters can be evaluated in a few clicks. In addition, users can share the platform and statistical results with collaborators and the public in different ways.

**Availability and Implementation:** The project page is https://github.com/mengchen18/omicsViewer

## Introduction

Thanks to the development of high throughput technologies such as next-generation sequencing and mass spectrometry, biology has transformed into a data-driven science exploiting the richness of omics data. The experimental design in the field has evolved from the simple “single-contrast” design to more complex designs, e.g., “multi-contrast” designs where multiple factors and their interactions are of interest. Some studies do not even have a clear pre-defined contrast, such as the Cancer Genome Atlas (TCGA)(“The Cancer Genome Atlas Program,” n.d.) and Clinical Proteomic Tumor Analysis Consortium (CPTAC)(Ellis et al. 2013). Usually, researchers measure multiple omics data over a large number of samples and use a wide range of phenotypic/clinical variables for data interpretation. The overall aim of these studies is to define cancer subtypes and elucidate biomarkers associated with clinical factors, such as responses to therapies, prognosis, etc. The biological interpretation of such omics data requires advanced statistical methods with fine-tuned parameters. However, thorough exploration of statistical results and evaluation of analysis parameters is often hampered by the high dimensionality of omics and phenotypic data. This process is even more challenging in projects with a highly collaborative environment where scientists from different backgrounds cooperate closely. Recently, the importance of interactive visualization of omics data was well recognized and several interactive tools have been developed, for example, the iSEE package (Rue-Albrecht et al. 2018) for the quick check of single or multiple omics data, MatrixQCvis (Naake and Huber 2021) for evaluating data quality and normalization methods, or VolcaNoseR (Goedhart and Luijsterburg 2020) for exploring volcano plots. In this work, we developed a general omics data visualization platform called “omicsViewer”. It is featured as interactive visualizations of advanced statistical results, links expression with phenotypic data, and performs downstream analyses on the fly in response to the selection of features and samples.

## Design concept

- **SummarizedExperiment-centric**. omicsViewer visualizes Bioconductor S4 class SummarizedExperiment (or ExpressionSet), which usually consists of three matrix-like objects: i) an expression matrix resulting from an omics study; ii) meta information of samples (referred to as phenotypic data); iii) meta information of features (referred to as feature data) of the expression matrix. The statistical results are usually stored in the phenotypic and feature data and will be visualized upon the selection by users.
- **Calculated in R and visualized in Shiny.** The object to be visualized by omicsViewer is prepared in R statistical environment (Computing and Others 2013). This guarantees maximum flexibility in the statistical analysis. Additionally, the package provides convenience functions for the most frequently used statistical methods in omics data analysis: t-test, correlation test, PCA, gene set annotation, and outlier analysis. Once the object is prepared, the visualization is powered by Shiny and Plotly, and no coding is needed. Therefore, users can easily explore statistical results interactively. For example, p-values and fold changes could be calculated and stored in the feature data. The shinyapp users can visualize features using the two variables in a volcano plot. Users may cluster samples and later use the cluster assignment to annotate or sort columns of a heatmap.
- **On-the-fly downstream analysis**. In omicsViewer different types of downstream analysis can be performed according to the selected samples (or features) and phenotypic variables. For more information, see “The user interface” section.
- **Extendable.** Advanced users can customize or create new analyses and visualizations to be integrated into the current scheme. Therefore, it is possible to include extra analyses for in-depth analysis of a particular field of omics studies, such as multi-omics integration, single-cell omics experiment, or post-translation modification experiments.
- **The flexibility of result delivery**. omicsViewer can be started inside an R environment, hosted on a shiny server to be accessed by the public or internal users, or be included in a standalone data package to be shared with collaborators or submitted to journals as supplementary data.

## The user interface

The layout of omicsViewer comprises two parts. The left panel versatile visualizes the expression matrix and meta information of features and samples, and it serves as a “feature/sample selector”. A subset of features and samples can be selected interactively and passed to the right panel. Then different types of downstream analysis, such as survival analysis and gene set analysis, can be performed based on the selected features or samples. Therefore, any variables useful for prioritizing features or samples should be stored in the feature and phenotypic data. The left panel uses four types of visualizations:

- A **Scatter Plot** is used when two quantitative variables are selected. For example, two principal components could be selected to visualize the similarity of samples from a given data set. In the two groups comparison, features can be visualized using a volcano plot, where fold changes are on the x-axis and logarithm transformed p-values on the y-axis (Figure 1A and B).
- **Beeswarm Plot** (Wilkinson 1999) is used to visualize a quantitative variable versus a categorical variable, e.g., group tumor samples according to the subtypes. Beeswarm plot, rather than boxplot, is used to guarantee that users can select every individual sample or feature.
- **Heatmap** is used to visualize the expression matrix. Here the rows are features and columns are samples. The interactive heatmap allows users to add annotation variables on the rows and columns as color bars or barplot, reorder rows and columns according to external variables, and zoom in on a specific area in the plot (Figure 1C).
- **Interactive Tables** are used to show all data within omicsViewer. Tables are automatically updated upon the selection of features or samples.

**Figure 1.**
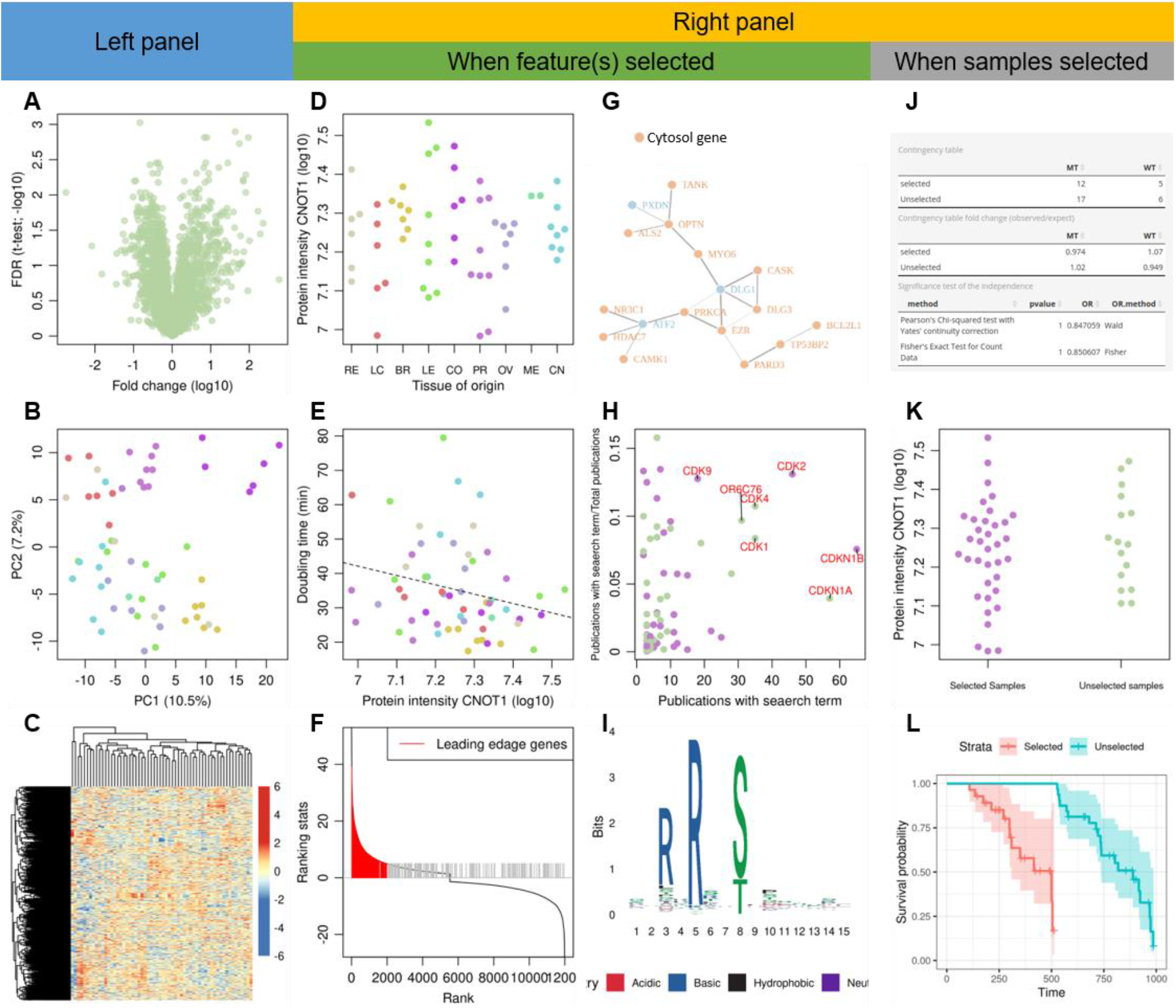
Example visualizations in omicsViewer. (A-C) Visualization used in the left panel. (A) Visualizing features in a volcano plot; (B) Visualizing samples using the first two principal components; (C) Heatmap shows the expression matrix. (D-I) Visualizations used in the right panel when features are selected. (D) Beeswarm plot shows the gene expression (x-axis) separated by a categorical variable (y-axis); (E) Scatter plot shows the correlation of two quantitative variables; (F) Visualization of fGSEA results: the rank statistic is sorted and shown as the solid curve, vertical gray bars shows genes annotated with a gene set term, red vertical lines mark the leading edge genes; (G) Example network returned by STRING database query; (H) Scatter plot shows the result returned by geneshot: points are genes, the x-axis shows the number of publication associated with a gene, and the y-axis shows the proportion of these publications also associated with the searched term. The most interesting genes are the ones on the top right corner. (I) An example sequence logo generated based on selected features. (J-L) Visualizations used in the right panel when samples are selected. (J) Examples tables show the results of chi-square and Fisher’s exact tests. (K) Beeswarm plot shows how a quantitative phenotypic variable differs between selected and unselected samples. (L) An example KM curve shows how the survival time is different between selected and unselected samples.

The analyses performed on the right panel when features are selected:

- **Significance test of mean** expression in two groups: If a single feature and a categorical phenotypic variable are selected, users can perform a Student t-test and Wilcox-test to test whether the mean expression of the feature is significantly different between any two groups in the phenotypic variable (Figure 1D).
- **Test of linear correlation:** If a single feature and a quantitative phenotypic variable are selected, the correlation of feature expression and the selected variable will be shown as a scatter plot. The significance test is performed using the correlation test (Figure 1E).
- **Over-representation analysis:** If users annotated features with gene set information, an over-representation analysis can be performed using Fisher’s exact test upon the selection of multiple features. The selected features are s the “feature of interest”, e.g., features most differentially expressed in two-conditions experimental design. All identified features are the background. In addition, annotations will be listed in a separate table for reference.
- **Fast Gene Set Enrichment Analysis (fGSEA)**: This analysis does not depend on the feature selection on the left panel. Instead, the fGSEA algorithm (Korotkevich et al. 2016) requires a ranking statistic, e.g., a principal component or fold-change comparison between two groups, to be specified on the right panel (Figure 1F).
- **Protein-Protein Interaction Networks:** Multiple selected features can be used to query the STRING database (Szklarczyk et al. 2021) for protein interaction analysis (Figure 1G). This function requires that features annotated with IDs acceptable for a STRING database query, e.g. gene symbol or UniProt IDs.
- **Geneshot**: Geneshot is a service that associates genes with biomedical terms (Lachmann et al. 2019) (Figure 1H). But unlike enrichment analysis, it allows free term searches. The associations between genes and terms are ranked, thus, it is sometimes more helpful to prioritize interesting candidate features.
- **Sequence logo:** If features are annotated with amino acid sequences and the base sequences of nucleic acids, a seqLogo (Wagih 2017) can be displayed based on the selected features (Figure 1I).

Analyses to be performed when a subset of samples are selected:

- When users select a categorical phenotypic variable, its association with the selected or unselected samples is tested using the Chi-square test and Fisher’s exact test (Figure 1J).
- When a quantitative phenotypic variable is selected, t-test and Wilcox-test are performed to examine whether the mean of the selected variable is significantly different between the selected and unselected samples (Figure 1K).
- When a variable for survival analysis is selected, Kaplan-Maier (KM) curves stratified by the selection of samples are shown. A log-rank test is used to test whether the selected and unselected samples have significantly different survival expectations (Figure 1L).

## Acknowledgments

The author thank Dr. Christina Ludwig and Dr. Isabella Straub for improving the graphical user interface and thank Dr. Christina Ludwig for reading and commenting on the manuscript.

## Reference

Computing, Rffs, and Others. 2013. “R: A Language and Environment for Statistical Computing.” Vienna: R Core Team. https://www.yumpu.com/en/document/view/6853895/r-a-language-and-environment-for-statistical-computing.

Ellis, Matthew J., Michael Gillette, Steven A. Carr, Amanda G. Paulovich, Richard D. Smith, Karin K. Rodland, R. Reid Townsend, et al. 2013. “Connecting Genomic Alterations to Cancer Biology with Proteomics: The NCI Clinical Proteomic Tumor Analysis Consortium.” Cancer Discovery 3 (10): 1108–12.

Goedhart, Joachim, and Martijn S. Luijsterburg. 2020. “VolcaNoseR Is a Web App for Creating, Exploring, Labeling and Sharing Volcano Plots.” Scientific Reports 10 (1): 20560.

Korotkevich, Gennady, Vladimir Sukhov, Nikolay Budin, Boris Shpak, Maxim N. Artyomov, and Alexey Sergushichev. 2016. “Fast Gene Set Enrichment Analysis.” BioRxiv. bioRxiv. https://doi.org/10.1101/060012.

Lachmann, Alexander, Brian M. Schilder, Megan L. Wojciechowicz, Denis Torre, Maxim V. Kuleshov, Alexandra B. Keenan, and Avi Ma’ayan. 2019. “Geneshot: Search Engine for Ranking Genes from Arbitrary Text Queries.” Nucleic Acids Research 47 (W1): W571–77.

Naake, Thomas, and Wolfgang Huber. 2021. “MatrixQCvis: Shiny-Based Interactive Data Quality Exploration for Omics Data.” Bioinformatics, November. https://doi.org/10.1093/bioinformatics/btab748.

Rue-Albrecht, Kevin, Federico Marini, Charlotte Soneson, and Aaron T. L. Lun. 2018. “ISEE: Interactive SummarizedExperiment Explorer.” F1000Research 7 (June): 741.

Szklarczyk, Damian, Annika L. Gable, Katerina C. Nastou, David Lyon, Rebecca Kirsch, Sampo Pyysalo, Nadezhda T. Doncheva, et al. 2021. “The STRING Database in 2021: Customizable Protein-Protein Networks, and Functional Characterization of User-Uploaded Gene/Measurement Sets.” Nucleic Acids Research 49 (D1): D605–12.

“The Cancer Genome Atlas Program.” n.d. The Cancer Genome Atlas Program. https://www.cancer.gov/tcga.

Wagih, Omar. 2017. “Ggseqlogo: A Versatile R Package for Drawing Sequence Logos.” Bioinformatics 33 (22): 3645–47.

Wilkinson, Leland. 1999. “Dot Plots.” The American Statistician 53 (3): 276–81.

